# Abnormal microstructural development of the cerebral cortex in neonates with congenital heart disease is associated with impaired cerebral oxygen delivery

**DOI:** 10.1101/332247

**Authors:** Christopher J. Kelly, Daan Christiaens, Dafnis Batalle, Antonios Makropoulos, Lucilio Cordero-Grande, Johannes K. Steinweg, Jonathan O’Muircheartaigh, Hammad Khan, Geraint Lee, Suresh Victor, Daniel C. Alexander, Hui Zhang, John Simpson, Joseph V. Hajnal, A. David Edwards, Mary A. Rutherford, Serena J. Counsell

**Author notes:** **Address for correspondence:** Professor Serena Counsell, Centre for the Developing Brain, School of Biomedical Engineering & Imaging Sciences, 1st Floor South Wing, St Thomas’ Hospital, London SE1 7EH.

## Abstract

**Background:** Abnormal macrostructural development of the cerebral cortex has been associated with hypoxia in infants with congenital heart disease (CHD). Animal studies have suggested that hypoxia results in cortical dysmaturation at the cellular level. New magnetic resonance imaging (MRI) techniques offer the potential to investigate the relationship between cerebral oxygen delivery and microstructural development of the cortex in newborn infants with CHD.

**Methods:** We measured macrostructural and microstructural properties of the cortex in 48 newborn infants with complex CHD and 48 age-matched healthy controls. Cortical volume and gyrification index were calculated from high resolution structural MRI. Neurite density and orientation dispersion indices were modelled using high angular resolution diffusion MRI. Cerebral oxygen delivery was estimated in infants with CHD using phase contrast MRI and pre-ductal pulse oximetry. We used tract-based spatial statistics to examine voxel-wise group differences in cortical microstructure.

**Results:** Microstructural development of the cortex was abnormal in 48 infants with CHD, with regions of increased fractional anisotropy (FA) and reduced orientation dispersion index (ODI) compared to 48 healthy controls, correcting for gestational age at birth and scan (FWE-corrected for multiple comparisons at P<0.05). Regions of reduced cortical ODI in infants with CHD were related to impaired cerebral oxygen delivery (R^2^=0.637, n=39). Cortical ODI was associated with gyrification index (R^2^=0.589, P<0.0001, n=48).

**Conclusions:** This study suggests that the primary component of cerebral cortex dysmaturation in CHD is impaired dendritic arborisation, which may underlie abnormal macrostructural findings reported in this population. The degree of impairment was related to cerebral oxygen delivery, supporting the hypothesis that maternal oxygen therapy may be beneficial in this population.

## Introduction

Congenital heart disease (CHD) is the most common congenital abnormality, affecting almost 1% of newborns^1^. Despite improvements in antenatal diagnosis, cardiac surgery and perioperative care, infants with CHD often experience neurological impairment across a range of developmental domains, both in early childhood and later in adult life^2,3^, making improvement of neurodevelopmental outcomes a major remaining challenge in the management of CHD. While early research focused on suspected surgical and perioperative factors, it now appears that a more complex set of biological factors may be responsible.

The detrimental effect of CHD on early brain development can be observed via a faltering trajectory of brain growth in the third trimester of pregnancy^4–7^, and a higher incidence of acquired brain lesions in newborn infants prior to cardiac surgery^8^. The developmental morphology of the cortex has become of increasing interest in CHD, with an ’immature cortical mantle’ first observed in autopsies of infants with hypoplastic left heart syndrome (HLHS)^9^. Reduced cortical folding has since been quantified *in vivo* using magnetic resonance imaging (MRI) in both fetal^6^ and pre-surgical neonatal populations^10–12^.Hypothesised contributory factors include reduced fetal cerebral oxygen delivery^5^ and cerebral metabolic substrate^13^, although precise cellular mechanisms remain unclear.

Linking physiological changes in CHD to brain development is assisted by four recent findings. Firstly, oxygen tension has been shown to regulate development of human cortical radial glial cells, with hypoxia exerting negative effects on gliogenesis by reducing the number of pre-oligodendrocytes while increasing the number of reactive astrocytes^14^. Secondly, hypoxia has been shown to reduce proliferation and neurogenesis in the subventricular zone of the piglet brain, accompanied by reduced cortical growth, with preliminary similarities found in the subventricular zone cytoarchitecture in human fetal autopsy specimens^15^. Thirdly, ascending aorta oxygen saturations have been found to be 10% lower in human fetuses with mixed CHD compared to healthy controls, with saturation measurements that correlated with fetal brain size^5^. Finally, microstructural maturation of the cortex, measured using both histology and diffusion anisotropy, has been demonstrated to occur in parallel with macrostructural development^16^. Taken together, these studies raise the hypothesis that macrostructural changes observed in CHD are the result of altered cortical microstructural development, which is in turn hindered by suboptimal oxygen tension during fetal life in CHD. Diffusion MRI, with newer multi-compartment models such as neurite orientation dispersion and density imaging (NODDI), provides measures that enable this hypothesis to be tested. Diffusion tensor imaging (DTI) metrics such as fractional anisotropy (FA) are non-specific and reflect many underlying parameters of brain tissue including neuronal density, fibre orientation dispersion, degree of myelination, free-water content, and axonal diameter^17^. The NODDI model aims to disentangle these different factors by separating the influence of neurite density and orientation dispersion from each other, and from partial volume with CSF, to provide distinct indices: orientation dispersion index (ODI), which captures the degree of dispersion of axonal fibre orientations (e.g. through fanning, bending, crossing) or dendrite orientations, and neurite density index (NDI), represented by the intracellular volume fraction^18^.

In this study, we aimed to use high angular resolution diffusion imaging (HARDI) and NODDI to test the hypothesis that reduced cerebral oxygen delivery in CHD is associated with impaired cortical microstructural development. We predicted that infants with complex CHD would exhibit higher cortical FA and lower ODI when compared to a group of healthy matched controls, and that infants with the lowest cerebral oxygen delivery would exhibit the most severe impairment of cortical microstructural development.

## Methods

The project was approved by the National Research Ethics Service West London committee (CHD: 07/H0707/105, Controls: 14/LO/1169) and informed written parental consent was obtained prior to imaging. All methods and experiments were performed in accordance with relevant guidelines and regulations.

## Participants

A prospective cohort of 54 infants with complex CHD expected to require surgery within one year was recruited from the Neonatal Intensive Care Unit at St Thomas’ Hospital, London. Six infants were excluded from the analysis: two infants with suspected coarctation were later assessed to have a normal circulation following postnatal ductus arteriosus closure; two infants were found to have focal arterial ischaemic stroke on MRI involving the cortex (both left middle cerebral artery stroke); one infant had uncertain gestation due to unknown date of last menstrual period and lack of ultrasound dating scan; one infant had incomplete diffusion data due to waking during the scan.

We therefore studied 48 infants with CHD, born at a median GA of 38.8 weeks (IQR 38.0 – 39.1). A control group of 48 healthy infants was matched by GA at birth and scan, recruited contemporaneously from the postnatal ward at St Thomas’ Hospital as part of the Developing Human Connectome Project^19^, and born at a median GA of 38.5 weeks (38.1 – 38.9). The median GA at scan was 39.1 weeks (IQR 38.6 – 39.7) for both the CHD group and control group.

## MR imaging

T1-weighted (T1w), T2-weighted (T2w), diffusion-weighted imaging (DWI), and phase contrast angiography (PCA) MR imaging was performed on a Philips Achieva 3 Tesla system (Best, The Netherlands) with a 32-channel neonatal head coil and neonatal positioning device^19^, situated on the neonatal intensive care unit at St Thomas’ Hospital, London. All examinations were supervised by a paediatrician experienced in MR imaging procedures. All infants were scanned in natural sleep without sedation. Pulse oximetry, respiratory rate, temperature and electrocardiography were monitored throughout. Ear protection comprised earplugs moulded from a silicone-based putty (President Putty, Coltene Whaledent, Mahwah, NJ, USA) placed in the external auditory meatus, neonatal earmuffs (MiniMuffs, Natus Medical Inc, San Carlos, CA, USA) and an acoustic hood positioned over the infant. All sequences included a 5 second initial slow ramp-up in acoustic noise to avoid eliciting a startle response.

T2w images were acquired using a multi-slice turbo spin echo sequence, acquired in two stacks of 2D slices (in sagittal and axial planes), using parameters: repetition time (TR): 12 s; echo time (TE): 156 ms, flip angle: 90°, slice thickness: 1.6 mm acquired with an overlap of 0.8 mm; in-plane resolution: 0.8×0.8 mm, scan time: 3:12 min per stack. The T1w volumetric magnetisation prepared rapid acquisition gradient echo acquisition parameters were as follows: TR: 11 ms, TE: 4.6 ms, TI: 714 ms, flip angle:9°, acquired voxel size: 0.8 × 0.8 × 0.8 mm, field of view (FOV): 145 × 145 × 108 mm, SENSE factor:1.2, scan time: 4:35 min. DWI with 300 directions was acquired using parameters: TR: 3.8 s, TE: 90 ms, multiband: 4; SENSE: 1.2; resolution: 1.5×1.5×3mm with 1.5mm slice overlap, diffusion gradient encoding: b=0 s/mm (n=20), b=400 s/mm (n=64), b=1000 s/mm (n=88), b=2600 s/mm (n=128) with interleaved phase encoding^20^. Quantitative flow imaging was performed using velocity-sensitised phase contrast imaging, with a single-slice T1-weighted fast field echo sequence. Scan parameters were: FOV: 100×100 mm, acquisition resolution: 0.6×0.6×4.0mm, TR: 6.4 ms, TE: 4.3 ms, flip angle: 10°, 20 repetitions, maximal encoding velocity (vENC): 140 cm/s, scan time: 71 s^21^.

## Structural and diffusion-weighted image reconstruction

T2w images were reconstructed using a dedicated neonatal motion correction algorithm. Retrospective motion-corrected reconstruction^22,23^ and integration of the information from both acquired orientations^24^ were used to obtain 0.8 mm isotropic T2w volumes with significantly reduced motion artefacts. Diffusion images were reconstructed following the scan using a dedicated multiband reconstruction method described previously^20^.

## Structural image processing

Motion-corrected T2w images were segmented into tissue type using an automated, neonatal-specific pipeline^25–27^, which was optimised for our acquisition parameters. Each tissue segmentation was manually inspected for accuracy using ITK-SNAP^28^, and minor corrections performed if necessary. Gyrification index was calculated as described previously^12^.

## Diffusion-weighted image processing

High angular resolution diffusion-weighted imaging (HARDI) data was reconstructed using a slice-to-volume motion correction technique that uses a bespoke spherical harmonics and radial decomposition (SHARD) of multi-shell diffusion data, together with outlier rejection, distortion and slice profile correction^29^. Data was first processed with image denoising^30^ and Gibbs ringing suppression^31^. A field map was estimated from b=0 images using FSL Topup^32^. Reconstruction was run for 10 iterations with Laplacian regularisation, using a SHARD decomposition of rank = 89 (allowing for spherical harmonics order lmax = 0, 4, 6, 8 for respective shells), with registration operating at a reduced rank = 15.

Non-brain tissue was removed using FSL BET^33^. DTI metrics FA and MD were calculated from b=0 and b=1000 DWI data using MRtrix3^34^. Neurite orientation dispersion and density imaging (NODDI) parameter maps were estimated using NODDI toolbox version 0.9^18^. We performed a Bayesian Information Criterion (BIC) comparison to compare the quality of the NODDI model fit for different intrinsic diffusivity values, which was found to be optimal at 2.0 × 10^−3^ mm^2^ s^−1^. This is consistent with previous NODDI studies in neonates^35–37^, with the higher value compared to adults (usually 1.7 × 10^−3^ mm^2^ s^−1^) likely reflecting the higher water content of the neonatal brain.

## Group template generation and image registration

A multivariate group template was generated from both T1w and T2w images, using symmetric diffeomorphic normalisation for multivariate neuroanatomy and a cross-correlation similarity metric^38^.Each subject’s diffusion data were rigidly registered to each subject’s T2w image using the MD map^39^. DTI and NODDI maps were then transformed into template space in a single step using concatenated linear and diffeomorphic transformations. Tissue segmentations from T2w images were also transformed into template space using nearest neighbour interpolation.

## Cortical analysis of microstructure

We used an approach for aligning cortical data from multiple subjects into a common space to provide voxel-wise spatial characterisation of FA, MD, NDI and ODI, as previously described^40,41^. A mean cortical map was produced by merging cortical grey matter segmentations that had been previously transformed into template space. This was then skeletonised to retain only a core of highly-probable cortical voxels, represented as a thin curved surface at the centre of the cortex. FA, MD, NDI and ODI measurements from each individual were projected onto the cortical skeleton by searching in a direction perpendicular to the cortical skeleton to identify voxels with the highest probability of being cortical.

## Calculation of cerebral oxygen delivery (CDO2)

For infants with CHD, we calculated their cerebral blood flow using a previously-described method^12^. Phase contrast angiography was acquired in a plane perpendicular to both internal carotids and basilar arteries at the level of the sphenoid bone^21^. Haemoglobin measurements were performed as part of routine clinical care, at a median of 3 days (IQR 0 – 5) prior to the scan. Arterial oxygen saturation (SaO_2_) was measured at the time of scan using a Masimo Radical-7 monitor (Masimo Corp, Irvine, CA) applied to the right hand.

Cerebral oxygen delivery (CDO_2_) was calculated using the following formulae^42^:

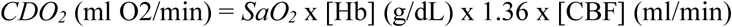

where 1.36 is the amount of oxygen bound per gram of haemoglobin at 1 atmosphere (Hüfner’s constant)^43^.

## Statistical analysis

Healthy infants were matched to the CHD group by both GA at birth and scan using an R implementation^44^ of the daisy algorithm^45^. In order to investigate the relationship between DWI metrics in the cortical grey matter and clinical factors, cross-subject voxel-wise statistical analysis was performed using FSL Randomise v2.9^41^. A general linear model (GLM) was used to assess group differences between diffusion measures of infants with CHD and healthy controls, including both GA at birth and scan as covariates in each model. The effect of CDO_2_ on cortical diffusion metrics was performed using a GLM that selected only infants with CHD who had a successful CDO2 measurement (n=39). All images were subject to family-wise error (FWE) correction for multiple comparisons following threshold-free cluster enhancement (TFCE)^46^ and are shown at P < 0.05. Linear regression was used to investigate the association between GI and diffusion metrics. To assess the relationship of GI and ODI independently of advancing brain maturity, GA at scan was included as a variable in the multiple linear regression model.

Categorical clinical variables were compared using Fisher’s exact tests. For continuous clinical variables, we determined medians and interquartile ranges, and compared groups using the Mann-Whitney U test. All analyses of clinical variables were performed using SPSS V24 (IBM, New York).

## Results

The analysis included 96 newborn infants: 48 infants with confirmed complex CHD scanned prior to surgery without evidence of arterial ischaemic stroke, and 48 age-matched healthy infants. Clinical characteristics of both groups are shown in Table 1. There were no significant differences in gestational age (GA) at birth, GA at scan and sex between groups. T1w, T2w, and DWI were acquired in all infants. PCA was acquired with acceptable quality in 81% of infants with CHD (n=39).

**Table 1.**
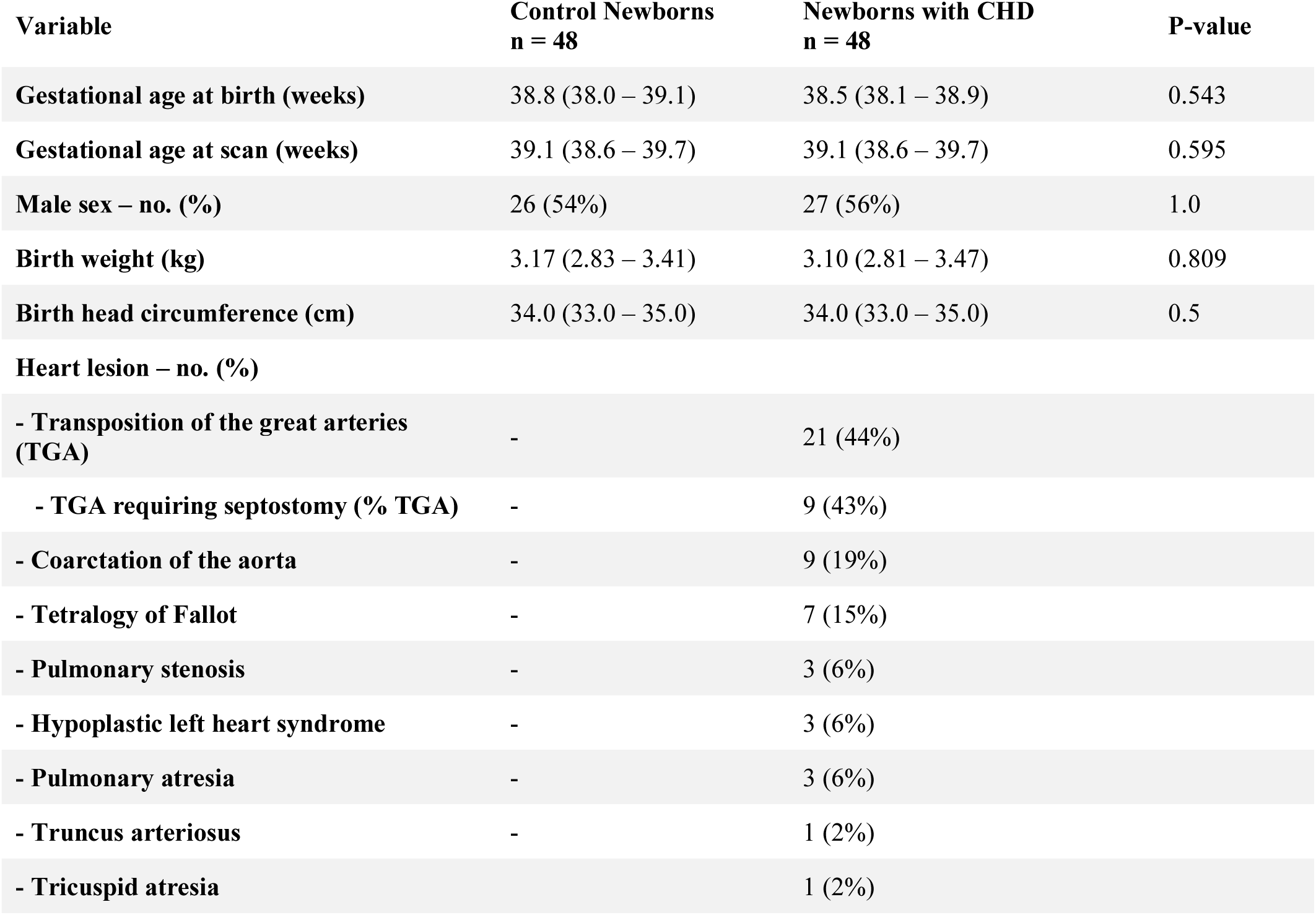
Clinical characteristics of the cohort. Values presented as median (interquartile range) unless otherwise stated. P-values calculated using Mann–Whitney U test for continuous data, and Fisher’s exact test for categorical variables. Septostomy in transposition of the great arteries (TGA) was performed prior to imaging in all cases.

### Cortical orientation dispersion index (ODI) is reduced in infants with complex CHD

Infants with CHD demonstrated widespread changes in cortical ODI, with the most significant reductions observed posteriorly in the posterior parietal cortex, insula cortex, cingulate cortex, primary motor cortex, supplementary motor area, and occipital regions (Figure 1a, corrected for GA at birth and scan). There were no regions where ODI was higher in infants with CHD. There were no differences in NDI between groups. GA at scan was positively associated with cortical ODI.

**Figure 1.**
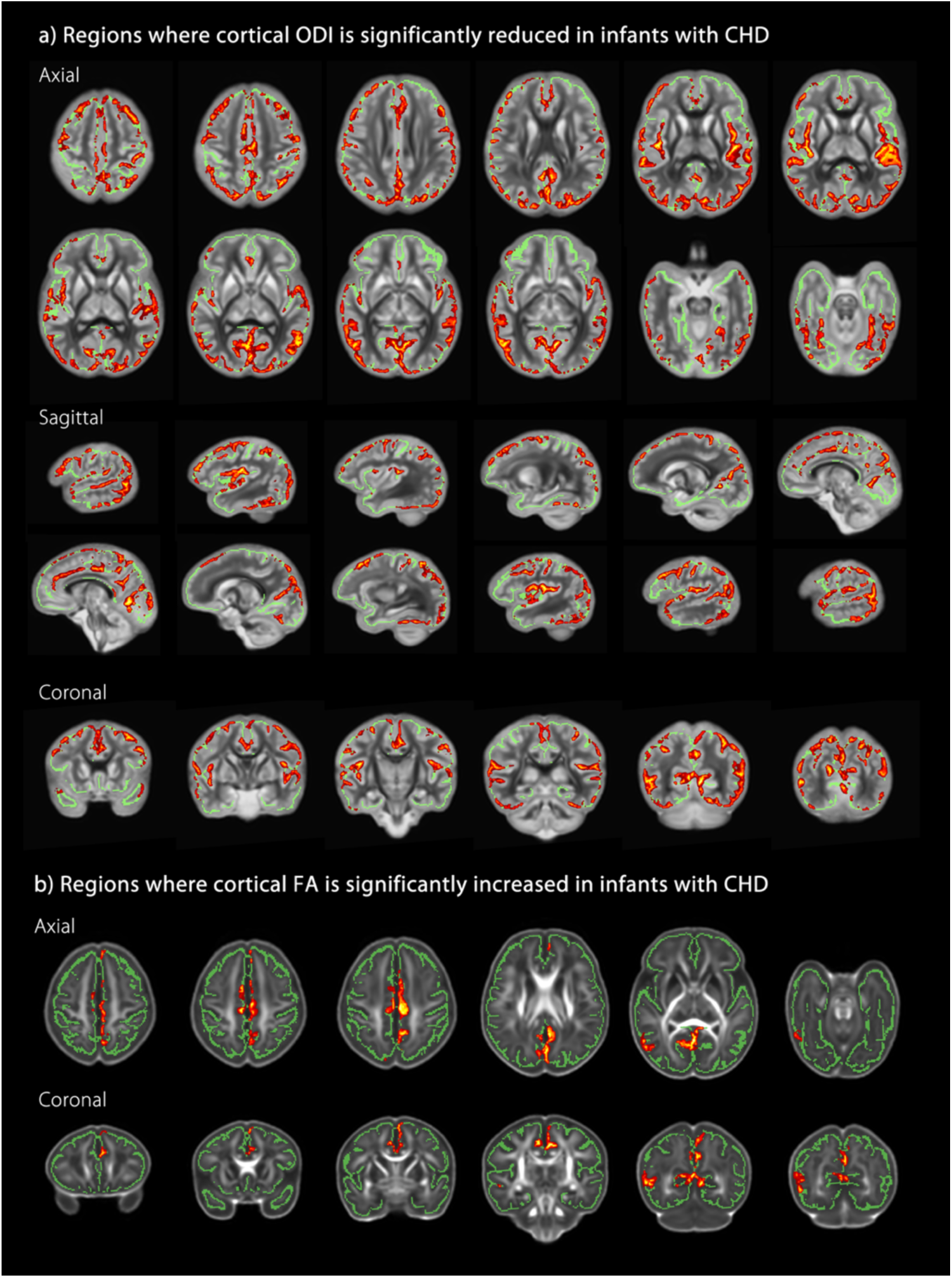
Infants with CHD (n=48) exhibit impaired microstructural development compared to healthy age-matched controls (n=48). **(A)** Regions where ODI is significantly reduced in infants with CHD, overlaid on the mean ODI template. **(B)** Regions where cortical FA is significantly increased in infants with CHD, overlaid on the mean FA template. Red-Yellow indicates P < 0.05 after FWE correction for multiple comparisons following TFCE. Results are shown overlaid on the mean cortical skeleton (green). Both analyses included GA at birth and at scan in the general linear model as covariates. Number of permutations was 10,000. Left–right orientation is according to radiological convention.

### Cortical fractional anisotropy (FA) is higher in infants with CHD

Cortical FA was higher in infants with CHD with effects seen in predominantly midline cortical structures (Figure 1b, corrected for GA at birth and scan). There were no regions where FA was lower in infants with CHD. There were no differences in mean diffusivity between groups. Cortical FA was negatively associated with GA at scan.

### Reduced cerebral oxygen delivery is associated with impaired cortical dispersion

Cerebral oxygen delivery (CDO_2_) at time of scan was positively associated with cortical ODI across many regions of the cortex (FWE-corrected for multiple comparisons, P < 0.05), with the most significant associations found in the bilateral temporal lobes, occipital lobes, cingulate cortex, and right insula cortex (Figure 2). To demonstrate this linear relationship, mean ODI data were extracted for each subject from significant voxels in the grey matter skeleton and plotted against CDO_2_ at time of scan (R^2^ = 0.637, Figure 3). There were no voxels with a negative association between the two variables. To assess the relative contribution of each component of CDO_2_, we repeated the analysis substituting CDO_2_ for either cerebral blood flow (CBF) or pre-ductal arterial saturation at time of scan. Considered alone, neither component demonstrated voxels that reached significance for either a positive or negative relationship with ODI. There were no significant associations between CDO_2_ and FA, MD or NDI.

**Figure 2.**
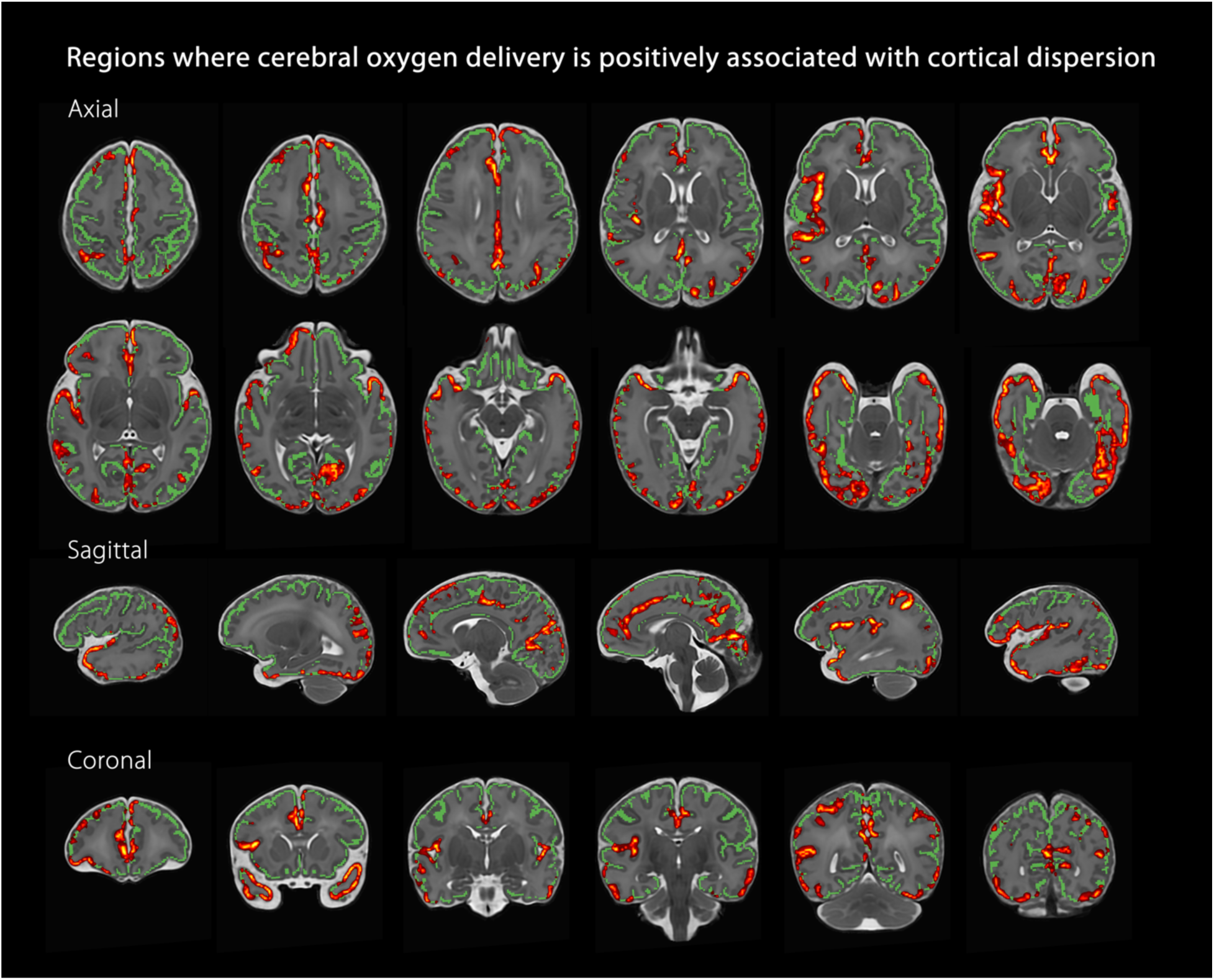
Regions where cerebral oxygen delivery is positively associated with cortical orientation dispersion in infants with CHD (n=39). Red-Yellow indicates P < 0.05 after FWE correction for multiple comparisons following TFCE. Results are shown overlaid on the group T2-weighted template and the mean cortical skeleton (green). Number of permutations was 10,000. Left–right orientation is according to radiological convention.

**Figure 3.**
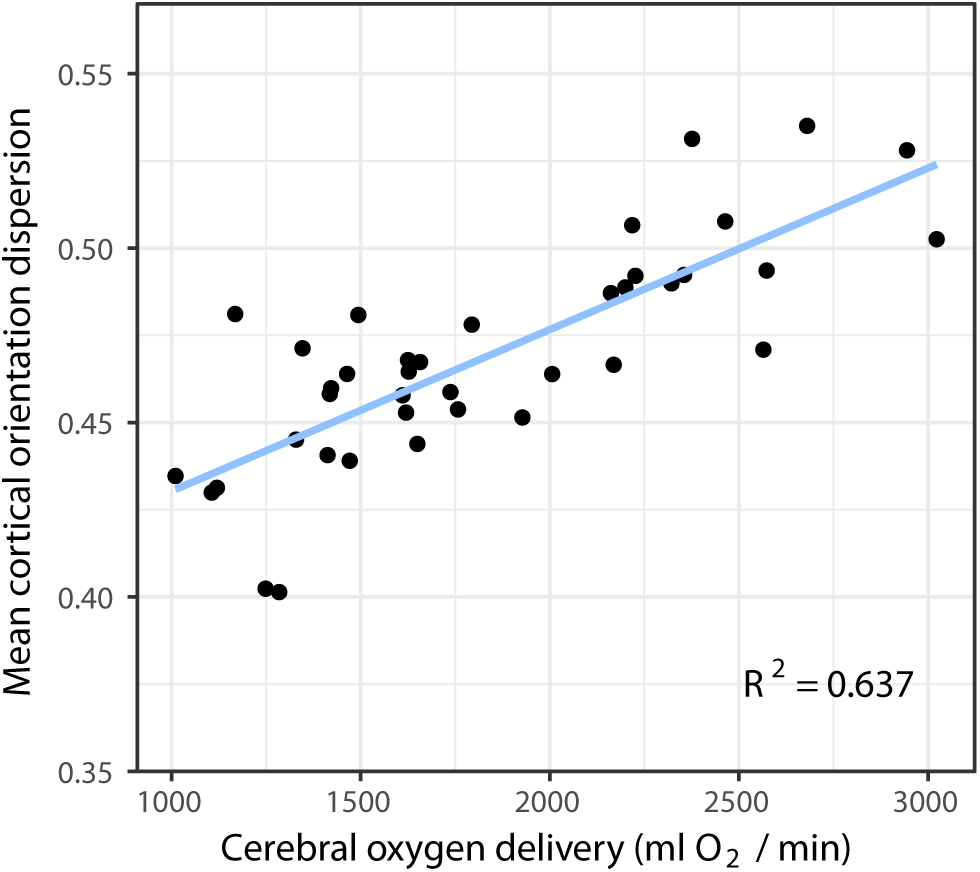
Visualisation of the linear relationship between CDO_2_ and cortical ODI within significant voxels from analysis displayed in Figure 2 (n=39). Mean ODI data were extracted for each subject from significant voxels in the statistic image after FWE-correction for multiple comparisons at P < 0.05.

### Relationship between cortical microstructure and macrostructure in CHD

We have previously reported reduced gyrification index (GI) in our cohort of infants with CHD^12^. However, the relationship between cortical microstructure and macrostructure has not been assessed in this group. We found that GI was significantly positively associated with mean cortical ODI (R^2^ = 0.589, P < 0.0001) and negatively with FA (R^2^ = 0.175, P = 0.003) (Figure 4). The linear relationship between GI and cortical ODI persisted following inclusion of GA at scan in the linear regression model (β = 0.642, P < 0.0001), but not for cortical FA (β = −0.304, P = 0.09). Cortical grey matter volume was significantly positively correlated with cortical ODI (R^2^ = 0.170, P = 0.004), but not with cortical FA. There was no relationship between total brain volume and microstructural measures (FA: R^2^ = 0.011, P = 0.49; ODI: R^2^ = 0.034, P = 0.207).

**Figure 4.**
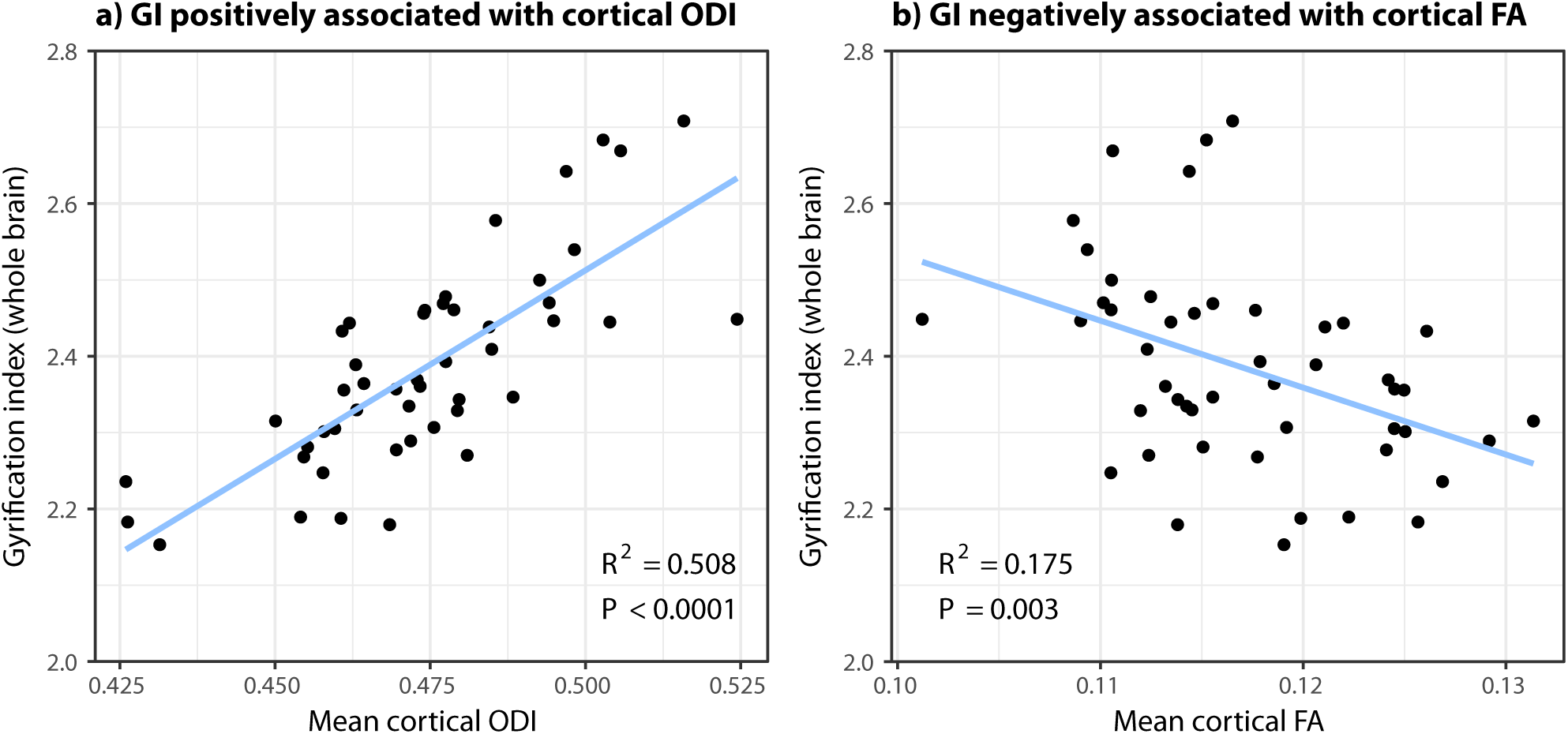
The linear relationship between gyrification index (GI) and cortical diffusion measures ODI and FA in newborn infants with CHD (n = 48). GI is (**a**) positively associated with cortical ODI and (**b**) negatively associated with cortical FA.

## Discussion

Long-term neurodevelopmental impairment is a major remaining challenge for infants with congenital heart disease, yet our understanding of the underlying biological substrate remains limited. Our study establishes that the microstructural development of the cerebral cortex in infants with CHD is abnormal in the newborn period compared to healthy controls, and importantly that the degree of impairment is related to reduced cerebral oxygen delivery. We speculate that hindered microstructural development underlies the abnormal macrostructural changes in brain development that have been observed through reduced birth head circumference^47^, smaller brain volumes^7^ and immature cortical folding^6,10–12,48^, and that strategies to optimise cerebral oxygenation in utero may offer the potential to ameliorate brain development in this population.

We found that cortical ODI was widely reduced in the CHD group, with associated but more sparsely distributed areas of higher FA. These findings suggest a hindered trajectory of normal brain development, with increased sensitivity to tissue changes using the more advanced NODDI model. As the brain matures in utero, cortical neurons migrate outwards towards the pial surface, populating the cortex^49^ and resulting in a highly directional, parallel, columnar microstructure. This can be observed with diffusion tensor imaging (DTI) as tensors with high FA, oriented radially to the cortical surface^50^. As the cortex matures, an increasingly dense and complex cytoarchitecture forms, with dendritic arborisation, glial proliferation, differentiation of radial glia and synapse formation^49,51,52^, associated with an observed increase in cortical ODI^35^. Increasing cytoarchitecture complexity also restricts water diffusion more evenly in all directions, with a consequent reduction in FA^40,50^. The link between diffusion imaging studies and underlying tissue biology is supported by prior evidence that NODDI-derived dispersion measures match their histological counterparts in adult postmortem specimens^53^, and by correlations found between maturation of dendritic arbors at the cellular level and loss of diffusion anisotropy with cortical development in the rhesus macaque^16^ and fetal sheep^54^.

While reduced cortical folding complexity in newborns with CHD has previously been reported in our cohort^12^, and others^11,55,56^, the link between cortical macrostructural and microstructural development in these infants has not been investigated previously. We found that gyrification index (GI) was positively associated with mean cortical ODI, independent of their association with increasing maturity. GI was also negatively but more weakly associated with mean cortical FA. Of interest, total brain volume was not associated with either FA or ODI. Taken together, these results suggest that macrostructural abnormalities observed in infants with CHD may be related to underlying impairments in dendritic arborisation.

Changes in cortical orientation dispersion were more pronounced posteriorly than frontally, which is consistent with previously described sequence of cortical development maturing earlier in the occipital cortex, and completing later in frontal regions^57–59^. Differences were also seen prominently in the region of the operculum, a region that has been repeatedly highlighted in infants with CHD. Findings of an ’open operculum’ with exposed insular cortex have been reported in CHD^48,60–63^, and has been associated with a poor outcome^64^.

Having established group differences between infants with CHD and healthy controls, we investigated the effect of CDO_2_ on development of cortical ODI. There was a widespread positive relationship between ODI and CDO_2_, supporting the hypothesis that impaired oxygen delivery to the developing brain may be associated with delayed cortical microstructural development. There was no relationship between cerebral blood flow alone and ODI, suggesting that impaired CDO_2_ rather than alternative proposed metabolic substrates^13^ may be most influential. These results support recent laboratory studies, with oxygen tension shown to regulate the development of human cortical radial glia cells. Moderate and severe levels of hypoxia exert negative effects on gliogenesis, mediated via reduced numbers of pre-oligodendrocytes and increased numbers of reactive astrocytes derived from cortical radial glia cells^14^. Equally, diminished subventricular zone neurogenesis as a result of chronic hypoxia may represent a cellular mechanism that underlies immature cortical development in the CHD population^15^. Taken together, these studies support the view that oxygenation of the developing brain is a crucial factor to optimise in order to restore the derailing trajectory of cortical development in this population. Future serial imaging to assess cortical development in CHD through infancy and early childhood would allow us to understand if microstructural differences observed in the newborn period are simply delayed with potential for catch-up in later childhood, or if such abnormalities are permanent and persist into childhood. Such work may offer insights into whether early surgical repair may help prevent further divergence, resulting in better long-term neurodevelopmental outcomes.

There were limitations to our study. Firstly, quantitative estimates obtained from microstructural studies are invariably model-dependent, exhibiting biases and limitations that are related to model assumptions. Despite this, NODDI indices have been shown to correlate with histological changes in neurite geometrical configuration^53^, and offers a valuable proxy to underlying biological changes in microstructure. Secondly, CDO_2_ was also measured in the postnatal period, while the most influential period on brain growth would have been in utero, and particularly during the third trimester. Despite this, we feel that postnatal CDO_2_ remains a useful surrogate for severity of cardiac circulatory compromise to date, taking into account both measures of cerebral blood flow and degree of hypoxia as a result of structural changes in congenital heart disease. Thirdly, the underlying genetic basis of CHD is becoming increasingly better-understood^65^, and may represent a key contributor not only to structural heart disease and associated impaired CDO_2_, but also to intrinsic abnormalities in microstructural development of the brain.

There are currently no validated neuroprotective therapies available for infants with CHD. Our demonstration that reduced CDO_2_ is associated with impaired cortical maturation in this population supports the development of strategies to optimise fetal CDO_2_. Strategies including the provision of supplemental oxygen to mothers during pregnancy may enable restoration of fetal cerebral oxygen tension to levels required to prevent or reverse abnormal corticogenesis. In addition, the use of newer microstructural measures such as ODI may provide a crucial leading indicator in the postnatal period to assess the impact of novel interventions on cortical development before child neurodevelopmental outcomes can be assessed at a later age.

## Acknowledgments

We are indebted to the families who supported this study. We thank staff from the St Thomas’ Neonatal Intensive Care Unit; the Evelina London Children’s Hospital Fetal and Paediatric Cardiology Departments; the Evelina London Paediatric Intensive Care Unit; the Centre for the Developing Brain at King’s College London; our research radiologists, including Sophie Arulkumaran, Kelly Pegoretti and Olivia Carney; our research radiographers, including Emer Hughes, Joanna Allsop, Ana Dos Santos Gomes and Elaine Green; Jiaying Zhang for assistance with diffusion modelling; and our neonatal scanning team including Katy Vecchiato, Julia Wurie, José Bueno Conde, Maryann Sharma, Beatriz Santamaria, Camilla OKeeffe and Jacqueline Brandon, whose energy and expertise made this study possible.

## Affiliations

Centre for the Developing Brain, School of Biomedical Engineering and Imaging Sciences, King’s College London, London (C.K., D.C., D.B., L.C-G.,J.K.S., J.OM.,S.V.,J.V.H.,A.D.E.,M.A.R.,S.J.C.);Biomedical Image Analysis Group, Department of Computing, Imperial College London, London (A.M.); Department of Forensic and Neurodevelopmental Sciences, King’s College London, Institute of Psychiatry, Psychology and Neuroscience, London (J.OM.), Department of Neuroimaging, King’s College London, Institute of Psychiatry, Psychology and Neuroscience, London (J.OM.); Neonatal Intensive Care Unit, St Thomas’ Hospital, London (H.K., G.L.); Department of Computer Science and Centre for Medical Image Computing, University College London, London (H.Z., D.C.A.,); Paediatric Cardiology Department, Evelina London Children’s Hospital, St Thomas’ Hospital, London (J.S.).

